# A novel truncating variant of GLI2 associated with Culler-Jones syndrome impairs Hedgehog signalling

**DOI:** 10.1101/365494

**Authors:** Fabiola Valenza, Davide Cittaro, Elia Stupka, Donatella Biancolini, Maria Grazia Patricelli, Dario Bonanomi, Dejan Lazarević

## Abstract

**Background:** GLI2 encodes for a transcription factor that controls the expression of several genes in the Hedgehog pathway. Mutations in GLI2 have been described as causative of a spectrum of clinical phenotypes, notably holoprosencephaly, hypopituitarism and postaxial polydactyl. Methods: In order to identify causative genetic variant, we performed exome sequencing of a trio from an Italian family with multiple affected individuals presenting clinical phenotypes in the Culler-Jones syndrome spectrum. We performed a series of assays, both in vitro and in ovo (Chicken model) to test the functional properties of GLI2 mutation.

**Results:** Here we report a novel deletion c.3493delC (p.P1167LfsX52) in the C-terminal activation domain of GLI2, and cell-based functional assays confirmed the pathogenicity of the identified variant and revealed a dominant-negative effect of mutant GLI2 on Hedgehog signalling.

**Conclusion:** Our results highlight the variable clinical manifestation of GLI2 mutations and emphasize the value of functional characterisation of novel gene variants to assist genetic counselling and diagnosis.

## Introduction

The Hedgehog (Hh) family of secreted morphogens control cell proliferation, differentiation and patterning during embryo development (Ingham, Nakano, & Seger, 2011). In humans, gene mutations in molecular components of the Hh pathway are associated with a number of congenital malformations or syndromes, characterized by abnormal development of brain structures, limbs and midline face, such as Holoprosencephaly (HPE), Greig cephalopolysyndactyly syndrome, Pallister-Hall Syndrome, Culler-Jones syndrome, and non-syndromic polydactylies (Bale, 2002). Besides its key roles in embryogenesis, Hh signaling remains active in some adult tissues, contributing to organ homeostasis and regeneration (Petrova & Joyner, 2014), whereas aberrant activation of the pathway occurs in several human cancers, including basal cell carcinoma and medulloblastoma (di Magliano & Hebrok, 2003).

All three vertebrate Hh ligands, Sonic hedgehog (*SHH*), Indian hedgehog and Desert hedgehog activate the same signal transduction pathway but operate in different tissues and organs. In the embryo, a concentration gradient of SHH, the most potent and best studied member of the family, specifies the identity of ventral neuron types along the entire rostral-caudal length of the central nervous system. SHH functions in the limb bud to define number, position and character of the digits (Tickle, 2006), and contributes to the development of the pituitary gland (Treier et al., 2001), cerebellum (Dahmane & Ruiz-i-Altaba, 1999; Wechsler-Reya & Scott, 1999), midbrain (Agarwala, Sanders, & Ragsdale, 2001), eye (Heavner & Pevny, 2012) and face (Xavier et al., 2016).

The effects of “canonical” Hh pathway are mediated by the GLI family of transcription factors (GLI1, GLI2 and GLI3), which control the expression of a number of target genes (Riobo & Manning, 2007). The three GLI proteins share a conserved C2H2-type zinc finger DNA-binding domain (Kinzler, Ruppert, Bigner, & Vogelstein, 1988). In addition, GLI2 and GLI3 possess an N-terminal repressor domain and a C-terminal activator domain, whereas GLI1 functions solely as an activator (Dai et al., 1999; Sasaki, Nishizaki, Hui, Nakafuku, & Kondoh, 1999). In the absence of Hh ligand, the membrane receptor Patched1 (PTCH1) inhibits the seven-transmembrane protein Smoothened (SMO) by preventing its access to the primary cilium. In this “off” state, GLI2 and GLI3 are retained in the cytoplasm by SUFU, a main negative regulator of the pathway, and undergo partial proteolytic processing that removes the C-terminal activation domain generating transcriptional repressor forms GLI3R and, to a lesser extent, GLI2R. Upon SHH binding, PTCH1 inhibition of SMO is relieved. As a result, GLI2 and GLI3 are converted into transcriptional activators (GLIA) that translocate into the nucleus to drive expression of target genes, including *GLI1* and *PTCH1*. Generally, GLI2A is the predominant activator of the pathway whereas GLI3R is the major transcriptional repressor, and their relative levels shape the SHH response (Briscoe & Thérond, 2013; Eggenschwiler & Anderson, 2007; Hui & Angers, 2011).

*GLI2* is a large and highly polymorphic gene, with a number of rare/family-specific heterozygous missense, non-sense, and frameshift mutations detected in individuals presenting with a spectrum of clinical phenotypes that include HPE, craniofacial abnormalities, polydactyly, panhypopituitarism, secondary hypogonadism or isolated growth hormone deficiency (Bear et al., 2014; França et al., 2010; Paulo et al., 2015; Roessler et al., 2003; 2005). The broad and variable range of clinical manifestations may have its origin in the bifunctional transcriptional activity of GLI2 and the complex regulatory feedbacks operating in the Hh signaling pathway, and points to possible modifying effects of additional genetic and environmental factors. Because it may be challenging to interpret the significance and impact of *GLI2* mutations, gene variants need to be functionally characterized to assess their pathogenicity and support genotype-phenotype correlation.

Here, we identify and functionally validate a novel truncating variant in the activation domain of GLI2 linked to a Culler-Jones syndrome phenotype characterized by hypopituitarism, polydactyly and facial dysmorphism in an Italian family.

## Material and Methods

### DNA Extraction and Sequencing

Genomic DNA (gDNA) was extracted from 800 μl of peripheral blood using the automated extractor Maxwell® 16 Research System (Promega, Madison, WI, USA); the concentration and high quality of gDNA (A260/280 1.8 to 2.0) was determined using a Nanodrop^TM^ Spectrophotometer 1000 (Thermo Fisher Scientific, Wilmington, DE, USA). Library preparation was performed using Illumina Nextera Expanded Rapid Caputer Enrichment Exome. Exome sequencing was carried out on Illumina HiSeq 2500 platform (Illumina, Inc. San Diego, CA, USA) using SBS chemistry. Libraries were sequenced in paired end mode, 101 nucleotides long each.

### Data Analysis

Reads were aligned to reference genome hg19 using bwa aln (Li & Durbin, 2010) (v 0.6.2). Variant calling was performed using GATK Unified Genotyper (Mckenna et al., 2010)(v.2.4.9) after Indel Realignment and VQSR according to GATK best practices (DePristo et al., 2011).

Variant effect prediction was performed using SnpEff v3.6 (Cingolani, Platts, Wang, Coon, & Nguyen, 2012) using GRCh37.34 genome version, subsequently variants were annotated to dbSNP v146 and to dbNSFP v2.4 (Liu, Jian, & Boerwinkle, 2013) using same suite.

The following chain of filters was applied to the variant set: GATK VQSLOD > 0, predicted change in coding sequence, segregation according to a dominant model, rarity in the population (COMMON = 0 in dbSNP build 137), predicted to be damaging according to SIFT (Kumar, Henikoff, & Ng, 2009) and Polyphen (Adzhubei et al., 2010). Genes were prioritized using the Phenolyzer platform (Yang, Robinson, & Wang, 2015).

### Expression Plasmids

The Human GLI2 cDNA clone was obtained from Addgene (pCS2-hGli2 #17648 (Roessler et al., 2005)) For N-terminal GFP tagging of GLI2, hGLI2 cDNA was inserted at the 3’-end of EGFP in the CMV expression vector pN1-EGFP (Clontech) using standard PCR-cloning. GeneArt Mutagenesis kit (Thermo) was used to introduce the deletion c.3493delC (p.P1167LfsX52) into GFP-tagged wild-type GLI2 expression construct using the following primers:

Forward 5’-CCAGCCAGGTGAAGCCTCCACCTTTCCTCAGGGCAACCTG-3’

Reverse 5’-CAGGTTGCCCTGAGGAAAGGTGGAGGCTTCACCTGGCTGG-3’

### Western Blotting

HEK293T cells (AD-293 cell line) were maintained in DMEM supplemented with 10% FBS, 1% LGlutamine, 1% Penicillin/Streptomycin. Cells transfected with the GFP-tagged constructs or pN1-EGFP control for 36 hrs using Lipofectamine 2000 (Thermo) were lysed on a nutator for 30 min at 4°C in lysis buffer (NaCl 150mM, EDTA 2mM, Tris-HCl pH7.5 50mM, Triton 1%) supplemented with protease and phosphatase inhibitor cocktails. After clarification by centrifugation at 13,000 g for 10 min at 4°C, 20μμg of total protein lysate were analyzed by SDS-PAGE followed by immunoblot with rabbit anti-GFP antibody (1:1000; Thermo) and rabbit anti-GAPDH (1:5000; Cell Signaling). Detection was performed by standard chemiluminescence with ECL Plus Western Blotting Substrate (Pierce/Thermo).

### Immunofluorence Assays

NIH-3T3 mouse fibroblasts were seeded on glass coverslips in DMEM supplemented with 10% FBS, 1% L-Glutamine, 1% Penicillin/Streptomycin for 24hrs, prior to treatment with 16nM SHH (Recombinant mouse SHH C25II N-Terminus, R&D Systems) in DMEM containing 0.5% FBS for 5 hrs. Cells were fixed for 20 min at RT with 4% PFA diluted in PBS and containing 4% sucrose, washed in PBS, stained with DAPI for 5 min and imaged with a 63x objective on a Leica TCS SP8 confocal microscope.

### Quantitative Real-Time PCR

NIH-3T3 cells were transfected in 12-well plate format with the indicated plasmids using Lipofectamine 2000 [1μg DNA:1.5μl Lipofectamine per well]. After 24 hrs, cells were treated with SHH (16nM) in DMEM containing 0.5% FBS for 40 hrs. 1 μg of total RNA extracted with Trizol (Thermo) was reverse transcribed into cDNA with M-MLV Reverse Transcriptase using random primers, and analyzed by SYBR green-based real-time quantitative PCR (SYBR Select Master Mix, Thermo) with the following primer sets:

*GAPDH* (F 5’-AATGTGTCCGTCGTGGATCTGA-3’; R 5’-AGAAGGTGGTGAAGCAGGCATC-3’),

*GLI1* (F 5’-TTATGGAGCAGCCAGAGAGA-3’; R 5’-ATTAACAAAGAAGCGGGCTC-3’)

*Ptch1* (F 5’-TGACAAAGCCGACTACATGC-3’; R 5’-AGAGCCCATCGAGTACGCT-3’)

### Luciferase Assays

NIH-3T3 were transfected in 24-well plate format using Lipofectamine 2000 with GFP-tagged wild-type GLI2, GLI2^MUT^ or GFP control plasmids along with a 8x-GLI-BS-Luc Firefly luciferase reporter construct containing eight repeats of the GLI binding sequence (Sasaki et al., 1999) and a plasmid expressing Renilla luciferase under the CMV promoter for normalization (Promega) in a 3:2:1 ratio (GLI2/mock GFP plasmids:8x-GLI-BS-Luc:Renilla Luc). [600ng total plasmid DNA:1μl Lipofectamine per well]. 24 hrs after transfection, cells were treated with SHH (16nM) in DMEM containing 0.5% FBS for 30 hrs and processed with Dual Luciferase Assay Kit (Promega) for measurement of Firefly and Renilla luciferase activities.

The same plasmids mixed at 2:1:1 ratio (GLI2/mock GFP plasmids:8x-GLI-BS-Luc:Renilla Luc) were electroporated “in-ovo” into neural progenitor cells in the neural tube of HH stage 13-14 chick embryos (Hamburger & Hamilton, 1992) using a square wave electroporator (BTX). After 24 hrs, the electroporated spinal cords were dissociated with trypsin supplemented with DNase (100U, Sigma) and cells were cultured for 1-2 hrs on PDL (100μg/ml)/laminin (1μg/ml)-coated 24-well plate in Neurobasal media containing B27 supplement (Gibco/Thermo Scientific), 2mM LGlutamine (Gibco/Thermo Scientific), 1% Penicillin/Streptomycin (Gibco/Thermo Scientific), 50μM Glutamic Acid (Sigma-Aldrich) (Bonanomi et al., 2012). Cells were then stimulated with SHH at the indicated concentrations for 24 hrs prior to harvesting and processing with Dual Luciferase Assay Kit (Promega).

## Results

### Clinical phenotype of the patients

Proband is a 6 year old female with neonatal panhypopituitarism (HP:0000871), prominent forehead (HP:0011220), thin upper lip vermilion (HP:0000219), downslanted palpebral fissures (HP:0000494), 2-3 finger syndactyly (HP:0001233), low-set ears (HP:0000369), single median maxillary incisor (HP:0006315), long philtrum (HP:0000343), bilateral postaxial hexadactyly (HP:0006136), choanal atresia (HP:0000453) and anterior pituitary agenesis (HP:0010626). Her father was diagnosed with hypopituitarism (HP:0040075) [hypothyroidism (HP:0000821), growth hormone deficiency (GHD) (HP:0000824)], unilateral hexadactyly (HP:0001162), and ectopic pituitary posterior lobe (HP:0011755). After further investigation, hexadactyly (HP:0001162) was described in the paternal grandfather, while GHD (HP:0000824) and hypopituitarism (HP:0040075) were identified in the paternal uncle who did not exhibit hand anomalies (Figure 1A).

**Figure 1.**
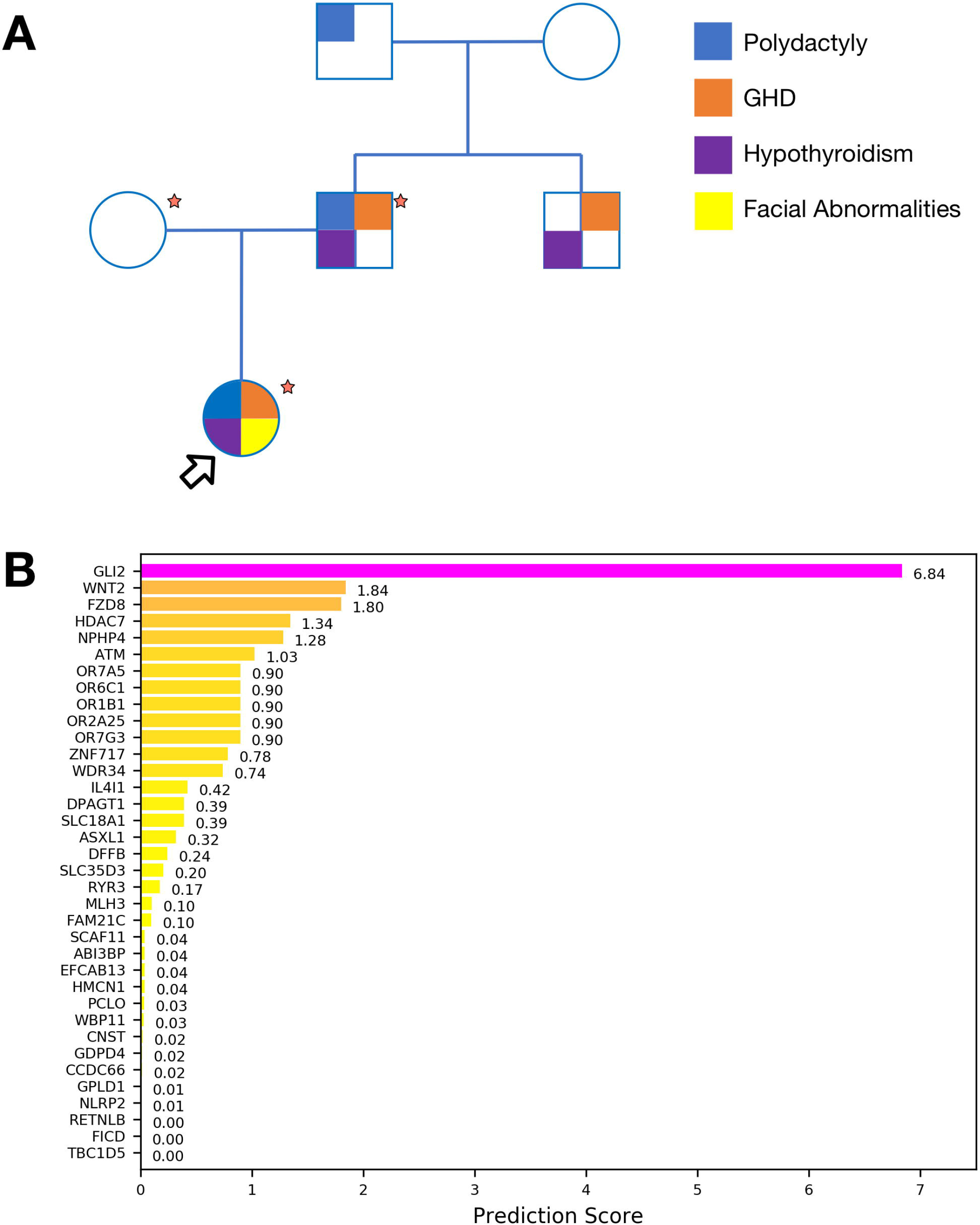
Exome Sequencing identifies *GLI2* as candidate disease gene: **(A)** Pedigree of the reported family. Main known phenotypes are mapped to individuals. Samples marked by a red star were available for Exome Sequencing. Proband is indicated with an arrow. **(B)** Bar plot representing prioritization scores obtained with Phenolyzer for genes identified from Exome Sequencing. *GLI2* can be effectively associated with the clinical phenotype.

The proband was previously tested negative for mutations in candidate disease-linked genes *POUF1*, *PROP1*, *HESX1, LHX3* and *GLI3*.

### Exome Sequencing identifies GLI2 as candidate disease gene

Exome sequencing was performed on the trio (Figure 1A) at average target coverage of 30x for each sample (Table S1). After variant calling, 130 rare or novel variants in 125 genes were found segregating according to a dominant model and affecting coding sequences. Once filtered for putative pathogenicity, 40 variants in 40 genes were retained (Table S2). Candidate genes were prioritized based on their association with the clinical phenotypes of the patients using Phenolyzer software. This analysis unambiguously identified *GLI2* as the top-scoring gene, while other candidates (*WDR34*, *DPAGT1*, *ASXL1*) already known to be associated with query phenotypes were ranked significantly lower (Figure 1B). Both the proband and her father were found to carry a novel heterozygous mutation caused by the deletion c.3493delC in the *GLI2* gene, leading to a frameshift and premature stop of translation at residue 1218 (p.P1167LfsX52).

### The mutation p.P1167LfsX52 converts GLI2 into a dominant-negative transcriptional repressor

The frameshift mutation c.3493delC in *GLI2* truncates the C-terminal portion of the transactivation domain required for transcriptional activity (p.P1167LfsX52, Figure 2A). To determine whether this truncation alters GLI2 function, we generated GFP-tagged constructs of either wild-type GLI2 or the p.P1167LfsX52 mutant (hereafter GLI2^MUT^) and examined their functional properties using cell-based assays. As predicted, GLI2^MUT^ revealed by western blotting in transfected HEK293 cells was smaller than the wild-type protein (~129kDa vs. 167kDa; ~156kDa vs. 194 kDa after GFP fusion) (Figure 2B). However, the subcellular distribution of GLI2^MUT^ expressed in NIH-3T3 mouse fibroblasts, which respond to SHH, was similar to that of wild-type GLI2: both proteins were found in the cytoplasm as well as the nucleus in untreated cells and accumulated within the nucleus following stimulation with SHH (Figure 2C-F’). Nevertheless, despite normal nuclear targeting, GLI2^MUT^ was unable to induce expression of the transcriptional targets of SHH signaling *GLI1* and *Ptch1*, whose mRNA levels were instead substantially higher in SHH-treated cells overexpressing wild-type GLI2 relative to mock-transfected controls (Figure 2G and H).

**Figure 2.**
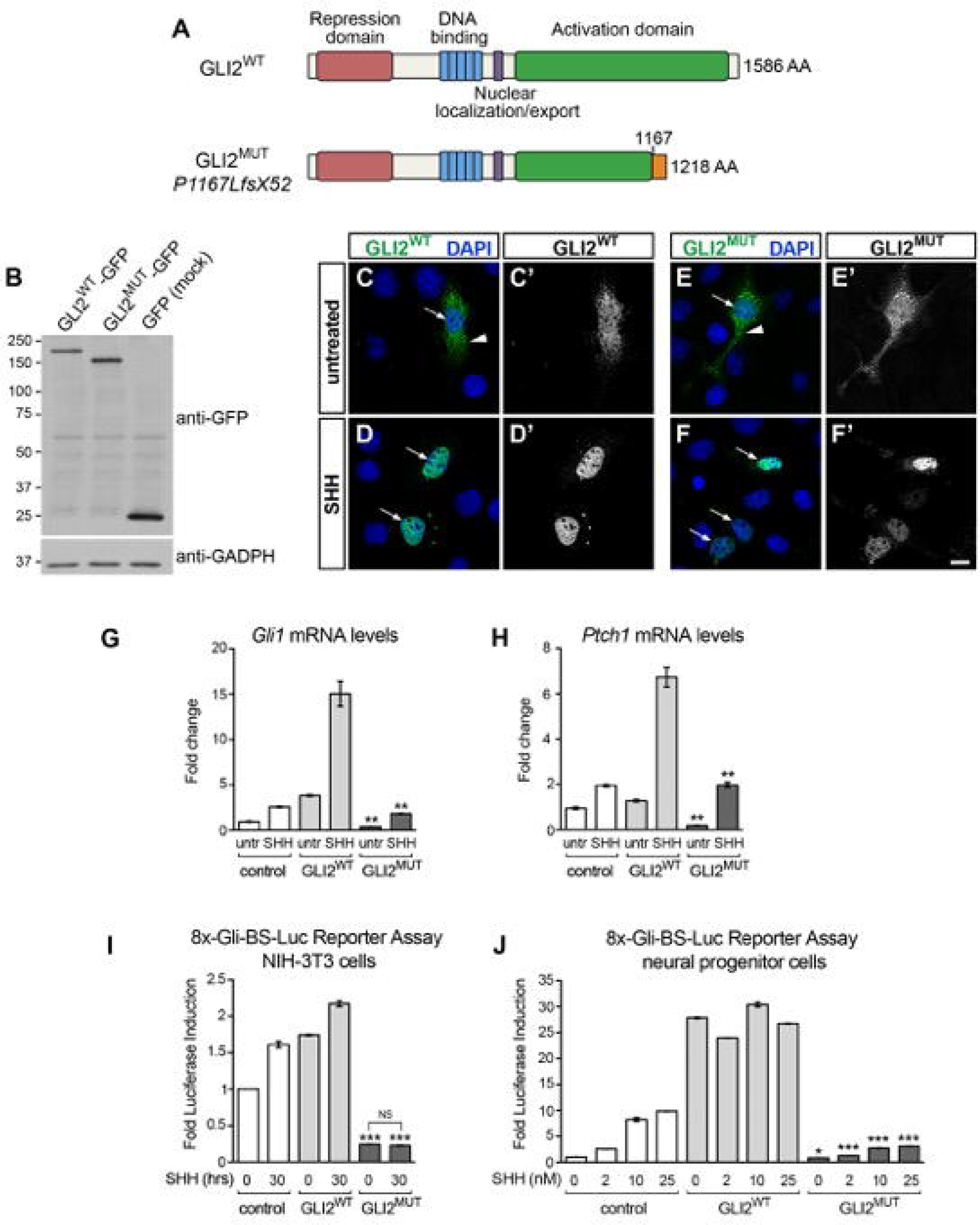
GLI2 mutant p.P1167LfsX52 lacks transcriptional activity and inhibits Hedgehog-GLI signaling: **(A)** Schematic of GLI2 wild-type and p.P1167LfsX52 mutant lacking the C-terminal region of the activation domain. Other functional motifs are intact, including the N-terminal repressor sequence, the zinc finger DNA-binding domain and nuclear localization signal. **(B)** Western blotting of total protein lysates of HEK293 cells transfected with plasmids expressing GFP-tagged wild-type human GLI2 (GLI2^WT^), p.P1167LfsX52 (GLI2^MUT^) or GFP (mock control) revealed with anti-GFP antibody. GADPH is a loading control. **(C-F’)** GFP-tagged GLI2^WT^ or GLI2^MUT^ visualized in transfected NIH-3T3 cells before or after stimulation with SHH (16nM) for 5 hrs. The GFP signal extracted from the corresponding merged images is show in C’-F’. Both proteins are found in the cytoplasm (arrowhead) and nucleus (arrow) in untreated cells and become primarily localized to the nucleus after stimulation. Scale bar, 10μm. (**G, H**) Levels of *GLI1* **(G)** and *PTCH1* **(H)** transcripts detected by quantitative-PCR in NIH-3T3 cells expressing GFP-tagged GLI2^WT^, GLI2^MUT^ or GFP control, before and after treatment with SHH (16nM) for 40 hrs. All conditions are normalized to untreated control cells (mean ± SEM, n=2). Unpaired *t*-test, (**) *p*<0.01 GLI2^MUT^ vs. GLI2^WT^ either untreated or SHH-treated matching conditions. (**I**) Luciferase-based reporter assay with GLI-responsive construct 8x-Gli-BS-Luc in NIH-3T3 cells transfected with GFP-tagged GLI2^WT^, GLI2^MUT^ or GFP control, before and after treatment with SHH (16nM) for 30 hrs. The expression levels of the reporter gene are measured by luciferase activity. All conditions are normalized to untreated control (mean ± SEM, n=2). Unpaired *t*-test (***), *p*<0.001 GLI2^MUT^ vs. control either untreated or SHH-treated matching conditions. (NS, non-significant) *p*=0.1264 GLI2^MUT^ untreated vs. treated. (**J**) Luciferase-based assay with 8x-Gli-BS-Luc reporter in chick spinal cord progenitor cells expressing GFP-tagged GLI2^WT^, GLI2^MUT^ or GFP control, treated with increasing doses of SHH for 24 hrs. All conditions are normalized to untreated control (mean ± SEM, n=2-4). Unpaired *t*-test, (*) *p*=0.0137 GLI2^MUT^ vs. control, untreated; (***) *p*<0.001 GLI2^MUT^ vs. control at corresponding SHH concentrations.

To directly investigate the effects of the p.P1167LfsX52 mutation on GLI2 transcriptional activity, we assessed the ability of GLI2^MUT^ to stimulate a GLI-dependent reporter construct (8xGliBS-Luc) in which a promoter containing tandem GLI responsive elements drives expression of firefly luciferase upon activation of the Hedgehog pathway (Sasaki et al., 1999). In NIH-3T3 cells, overexpression of wild-type GLI2 increased reporter activity in a ligand-independent manner to an extent comparable to control cells treated with SHH. Conversely, the basal levels of reporter activity were significantly lower in cells transfected with GLI2^MUT^ and did not show the expected increase after SHH stimulation (Figure 2I).

A complementary set of experiments was conducted in primary cultures of neural progenitor cells derived from the chick embryo neural tube electroporated with either wild-type or mutant GLI2 together with the 8xGliBS-Luc reporter. Spinal cord progenitors depend on graded SHH signaling to acquire class-specific molecular identities during embryo development (Briscoe, Pierani, Jessell, & Ericson, 2000) and exhibit reliable dose-dependent responsiveness to SHH in culture (Figure 2J, control). Electroporation of wild-type GLI2 led to robust induction of reporter activity independent of SHH stimulation, whereas GLI2^MUT^ caused a considerable reduction in luciferase levels compared to control cells at all ligand concentrations tested, indicating that the mutant protein suppresses transcription mediated by endogenous GLI factors (Figure 2J).

In conclusion, the truncated mutant p.P1167LfsX52 functions as a transcriptional repressor that exerts dominant-negative effects on GLI-dependent gene expression.

### Discussion and Conclusions

This study expands the spectrum of *GLI2* mutations reporting a novel heterozygous pathogenic variant (p.P1167LfsX52) that results in autosomal-dominant developmental abnormalities including polydactyly, hypopituitarism, GHD and hypothyroidism.

Functional studies based on cellular assays demonstrated that the frameshift mutation p.P1167LfsX52 truncates the C-terminal transactivation domain of GLI2 generating a transcriptional-repressor form that retains the ability to translocate into the nucleus in response to SHH but exhibits dominant-negative activity. As a result, we observed a significant inhibition of GLI reporter levels in cells expressing GLI2 p.P1167LfsX52, indicating that the activity of wild-type GLI proteins is suppressed by the mutant variant. Dominant-negative activity was reported for other pathogenic variants of *GLI2* with deletions in the activation domain (Roessler et al., 2005). The inhibitory effect was found to require integrity of the DNA-binding and amino-terminal transcriptional repressor domains (Flemming et al., 2013; Roessler et al., 2005), which are intact in GLI2 p.P1167LfsX52. To inhibit positive GLI function, C-terminally truncated variants may compete with and displace wild-type GLI2 from target sites and/or form inactive complexes with the activating forms.

While *GLI2* mutations were originally identified in patients with HPE and midline abnormalities, (Roessler et al., 2003; 2005), more recently it became clear that *GLI2* variants are often associated with polydactyly, pituitary deficiency and subtle midfacial facial phenotypes rather than patent HPE (Bear et al., 2014; França et al., 2010; Rahimov, Ribeiro, de Miranda, Richieri-Costa, & Murray, 2006) (Kordaß, Schröder, Elbracht, Soellner, & Eggermann, 2015). The fact that frank HPE is rare in patients with *GLI2* mutations, in contrast to those with *SHH* variants, has suggested that other GLI proteins (GLI1, GLI3) might function redundantly to compensate in part for GLI2 deficiency, in line with studies in compound mutant mice (Arnhold, França, Carvalho, Mendonca, & Jorge, 2015; Park et al., 2000; Sasaki et al., 1999). Likewise, *GLI2*-null mice display normal limb patterning unless *GLI1* is also ablated, and *GLI2/GLI3* double heterozygous mice have a more severe polydactyly than *GLI3* mutants (Park et al., 2000). Interestingly, polydactyly is generally present in patients with more severe *GLI2* variants, including those that disrupt the zinc-finger and transactivation domains (Arnhold et al., 2015). Specifically, individuals with mutations predicted to result in protein truncation are significantly more likely to present both polydactyly and pituitary insufficiency compared to those with non-truncating variants (Bear et al., 2014)

There is striking variability in the phenotypic outcomes of GLI2 mutations even within the same family tree (França et al., 2010; Paulo et al., 2015; Rahimov et al., 2006; Roessler et al., 2003; 2005), ranging from unaffected carriers to patients with craniofacial abnormalities, pituitary phenotypes and polydactyly, either isolated or in combination. In the pedigree examined in this study, hormone deficiencies, but not hand and facial anomalies, were present in all individuals carrying mutant *GLI2*, in support of the recommendation to consider this gene as a primary candidate to screen after endocrinology testing has revealed pituitary insufficiency even in the absence of polydactyly (Bear & Solomon, 2015). Incomplete penetrance and variable phenotypes, as also reported in patients with autosomal dominant mutations in other loci linked to HPE or pituitary deficiencies (e.g., *SHH*, *SIX3*, *OTX2*, *HESX1*), suggest the contribution of additional genetic variants, epigenetic changes and environmental factors. Despite the pathogenetic role of *GLI2* mutations has already been described, a small number of rare and damaging variants in the same gene can be found in public data from non-dysmorphic individuals such as the Exome Aggregation Consortium (Lek et al., 2016) or Exome Variant Server (Fu et al., 2012). Therefore, systematic functional validation of putative pathogenic variants would be valuable for genetic counseling and patient screening.

## Acknowledgements

This work was supported by the Career Development Award of the Giovanni Armenise-Harvard Foundation and by the European Research Council under the European Union’s Seventh Framework Programme (FP7/2007-2013)/ERC-StG 335590 NEVAI to D. Bonanomi; It was supported by Italian Ministry of Health (5 per mille).

## Conflict of interest

The authors declare that they have no competing interests.

## Patient consent

Obtained.

## Ethical approval

This study was approved by the Institutional Ethical Review Committee, San Raffaele Hospital, Milan, Italy (prot. RARE-DISEASE)

